# Impacts of landscape composition, marginality, and climatic stability on the patterns of endemism of Cerrado woody plants

**DOI:** 10.1101/362475

**Authors:** João de Deus Vidal Júnior, Anete Pereira de Souza, Ingrid Koch

## Abstract

**Aim:** Although various theories have been proposed to explain the outstanding endemism of plants in the Cerrado, four hypotheses about the mechanisms of diversification and distribution are most supported: (1) plateau/valley, (2) stable/unstable climate, (3) core/peripheral distribution, and (4) soil fertility. The first argues that plateaus harbor more ancient lineages than valleys and therefore presents higher endemism. The second theory suggests that climatic stable environments maintained more paleoendemic species. The third scenario attributes the distribution of endemism to gradients of conditions available to locally adapted species and predicts higher endemism in nuclear than in marginal areas. The last theory suggests that lower fertility soils account for higher endemism due to the habitat specialization of its species. We compared endemism patterns with the predictions of each theory to discuss their importance.

**Location:** Brazil.

**Time period:** Quaternary.

**Major taxa studied:** Angiosperms.

**Methods:** We mapped the endemism using records of 311 plant species of the Cerrado and applied spatial analysis and distribution models to summarize the importance of each predictor of endemism.

**Results:** We identified 28 areas in which the higher endemism of Cerrado plants were concentrated and presented a map of its distribution. We found correlations among endemism, climate stability, elevation, and marginality, which supported the plateau/valley, core/peripheral, and stable/unstable hypotheses. No association between soil fertility and endemism was detected. We propose that plateaus are more stable climatic environments, and this characteristic along with their elevation and centrality are predictive of endemism.

**Main conclusions:** We concluded that most of the endemism is concentrated in overlapping areas of stability of species, which are concentrated in higher elevation central regions. Soil fertility was not linked to endemism. We recommend that central plateaus in the Cerrado require special attention in conservation to optimize the protection of endemic species in the biome.

## INTRODUCTION

The Cerrado is the largest savanna and the second largest ecoregion in South America (Olson et al., 2001), and it has a complex and unique biogeographic history. This ecosystem harbors more plant species than any other savanna in the world (> 7000 species; Mendonça et al., 1998), with over 44% of its flora being endemic, thus making it the richest tropical savanna in the world in this regard (Klink & Machado, 2005). The Cerrado landscape is divided into ancient plateaus, which are large extension areas with relatively high elevations (500–1700 m), and peripheral valleys, which are surrounding plateaus with lower elevation (100–500 m) (Cole, 1986). Many different vegetation physiognomies are found in the Cerrado, and each presents a unique biota and different biogeographic history, including savanna woodlands, gallery forests, riverine forests, dry forests, marshlands, and enclaves with species from adjacent biomes (Cole, 1986; da Silva & Bates, 2002).

Various theories have been proposed to explain the high degree of endemism found for several taxa; however, no prevailing process acting on the biogeographic history of this domain has emerged. The endemism within a region is positively influenced by the speciation rate and decreases with local extinction and immigration rates (Chen & He, 2009), which are processes intrinsically linked to the biogeographical history. Currently, the most widely discussed hypotheses for the mechanisms underlying species diversification in the Cerrado include (1) the geographic compartmentalization of the landscape in plateaus and valleys of the Late Miocene, which may have driven of allopatric speciation between disjoint landscape units; (2) variability in climatic stability during the Pleistocene along regions within the Cerrado, which may have promoted differential maintenance of diversity in regions with stable vs. unstable climates; (3) the gradient of optimal environmental conditions between core and periphery of the Cerrado, which may have promoted lower extinction rates in nuclear regions and favored the maintenance of endemism in core areas of this domain; and (4) the unequal distribution of patches of low fertility soils, which may have concentrated the occurrence of endemism rich communities compared with patches of high fertility soils. Although these hypotheses are not mutually exclusive, the relative importance of these theories remains a major focus of debate because patterns observed in distinct taxa seem to support different scenarios (e.g., Turchetto-Zolet, Pinheiro, Salgueiro, & Palma-Silva, 2013).

Considering the four mentioned hypotheses, the first attributes differential endemism across the Cerrado to the older geological origin of plateau formations, which date from the Late Tertiary, and valleys to recent erosive processes (Ab’Sáber, 1983; Cole, 1986; da Silva, 1997; Werneck, 2011). Given this scenario, ancient geomorphological processes acting on these environments might have promoted the isolation of older lineages in the highlands.

Consequently, the recently formed valleys were gradually occupied by neoendemic species (Cole, 1986). Therefore, according to this hypothesis, plant communities in higher elevation regions are expected to present more endemic species than those occurring in adjacent depressions given the older establishment and relative isolation of higher regions (Cole, 1986; Werneck, 2011). This model is consistent with other previously reported endemism patterns for plants of the genus *Mimosa (*Simon & Proença, 2000), squamate lizards (Nogueira, Ribeiro, & Costa, 2011), and several birds (da Silva, 1997).

In addition to the geomorphological processes during the Late Miocene, the Pleistocene climate is proposed as a key driver of taxa diversification in the Cerrado (Werneck, Nogueira, Colli, Sites, & Costa, 2012; Santos, Nogueira, & Giugliano, 2014; Ribeiro *et al*., 2016; Bueno et al., 2017; Arruda, Schaefer, Fonseca, Solar, & Fernandes-Filho, 2018; Costa et al., 2018). In general, climatically stable regions are considered good predictors of diversity at both interspecific (Jansson, 2003; Werneck, 2011) and intraspecific levels (Carnaval & Moritz, 2008; Carnaval, Hickerson, Haddad, Rodrigues, & Moritz, 2009; Correa Ribeiro, Lemos-Filho, J.P., de Oliveira Buzatti, R.S., Lovato, M.B. & Heuertz, 2016; Buzatti, Lemos-Filho, Bueno, & Lovato, 2017; Lima *et al*. 2017; Collevatti *et al*. 2018). Areas presenting less stable climatic conditions are more prone to local extinction than regions with more stable climates due to the reduced population sizes and, consequently, greater importance of genetic drift and inbreeding (Harrison & Noss, 2017). Conversely, populations in climatically stable regions have been demonstrated to show higher demographic stability due to their more highly effective population sizes compared with that of climatically unstable areas (Correa Ribeiro *et al*. 2016; Buzatti *et al*. 2017; Lima *et al*. 2017; Collevatti *et al*. 2018). According to this hypothesis, overlapping stable areas across the distribution of multiple taxa may potentially constitute higher endemism regions as demonstrated for the Atlantic Rainforest (Carnaval & Moritz, 2008).

The third hypothesis suggests that core regions of the Cerrado have historically presented relatively more optimal conditions for the maintenance of regionally adapted populations of its species, thus supporting the diversification of endemic species through time (Soule, 1973; Eckert, Samis, & Lougheed, 2008). In this scenario, nuclear regions of the Cerrado are expected to show higher levels of endemism, whereas communities in peripheral regions are more prone to undergo stochastic events of local extinction, given the exposure of populations of species to sub-optimal conditions in comparison with nuclear regions (Soule, 1973). Moreover, the floristic composition of marginal environments may be under different environmental filters than adjacent vegetation types, which can result in unique floristic compositions and endemism levels (Neves et al., 2017).

The distribution of fertile soils represents an important factor that determines the vegetation structure and species distribution in the Cerrado (Dantas & Batalha 2011, Reatto *et al.* 2008, Franco, Rossatto, de Carvalho Ramos Silva, & da Silva Ferreira, 2014). Savanna species, for example, are typically established in acidic, sandy and well-drained soils with high aluminum and low calcium, magnesium, phosphorus, and nitrogen availability (Furley & Ratter, 1988; Haridasan, 2008). Low fertility soil patches or rocky outcrops, however, usually harbor more narrowly endemic species because these areas impose strong physiological selective pressures that demand specific adaptations (Médail & Verlaque, 1997; Beard, Chapman, & Gioia, 2000; Casazza, Barberis, & Minuto, 2005). In addition, these environments present low vegetation cover that may limit biotic interactions (Imbert et al., 2012). Thus, following this hypothesis, areas presenting low soil fertility are expected to present higher levels of endemism as a consequence of the habitat specialization of the species (Beard et al., 2000).

Multiple species niche models are useful for detecting general distribution patterns, which are important for assembling large-scale scenarios. Large-scale hypothesis testing for the identification of refugia has been conducted for several regions around the world, with important biogeographic outcomes (Thuiller, 2004; Svenning, Normand, & Kageyama, 2008). Mostly, single species patterns or simulated dataset models have been applied in past distributional reconstructions for the Cerrado (Collevatti et al., 2012; Werneck et al., 2012; Lima, Telles, Chaves, & Lima-Ribeiro, 2017; Arruda et al., 2018), whereas multiple species approaches are less common (de Siqueira & Peterson, 2003; Bonaccorso, Koch, & Peterson 2006; Bueno et al., 2017). These studies addressed future climatic changes based on a conservation biology approach (de Siqueira & Peterson, 2003), simulated the history of Amazonian forest fragmentation and the potential occupation of open areas by Cerrado species (Bonaccorso, Koch, & Peterson 2006), or reconstructed the distribution of woody savanna vegetation under past scenarios (Bueno et al., 2017). Most of these studies identified important relationships between landscape composition and shifts in the distribution of the savanna during glaciations. An additional possibility for testing hypotheses on the abiotic determinants of the endemism distribution along the Cerrado that has not yet been explored is the implementation of niche reconstructions and climatic projections to multiple species range shifts. Here, we compared the levels of endemism of the woody flora of the Cerrado domain with the three scenarios proposed to explain its diversification. We tested whether higher elevations, lower soil fertility, central region location, or climatically stable areas show higher levels of endemism. In addition, we modeled real species and simulated occurrence data of woody plants from valleys and plateaus to identify regions of stability and reconstruct potential retraction/expansion of environmental conditions occurring in these landscapes during the climatic shifts of the Quaternary.

## METHODS

### Species occurrence data and endemism calculation

To generate the maps of endemism for woody species of the Cerrado, we gathered occurrence data of 311 species from savannas, seasonal forests, and gallery forests (Fig. 1a, 1b, and 1c). We elaborated the species list for the savanna, gallery and seasonal forests based on Ribeiro and Walter (1998), the NeoTropTree database (Oliveira-Filho, 2017), and the SDTF catalog proposed by Prado and Gibbs (1993). We excluded species present in multiple vegetation types to reduce the bias of widespread species in models. We retrieved species occurrence data from GBIF using the R package “rgbif” (Chamberlain, Ram, Barve, & Mcglinn, 2016). To ensure the quality of the data, we adopted search parameters that restrict data without original coordinates and with spatial issues. We also discarded occurrences with less than three decimal digits of coordinate precision to avoid spatially biased data. We manually checked points by plotting individual maps for each species and discarding putative outliers. A full list of material data sources is presented in Appendix 2.

**Figure 1.**
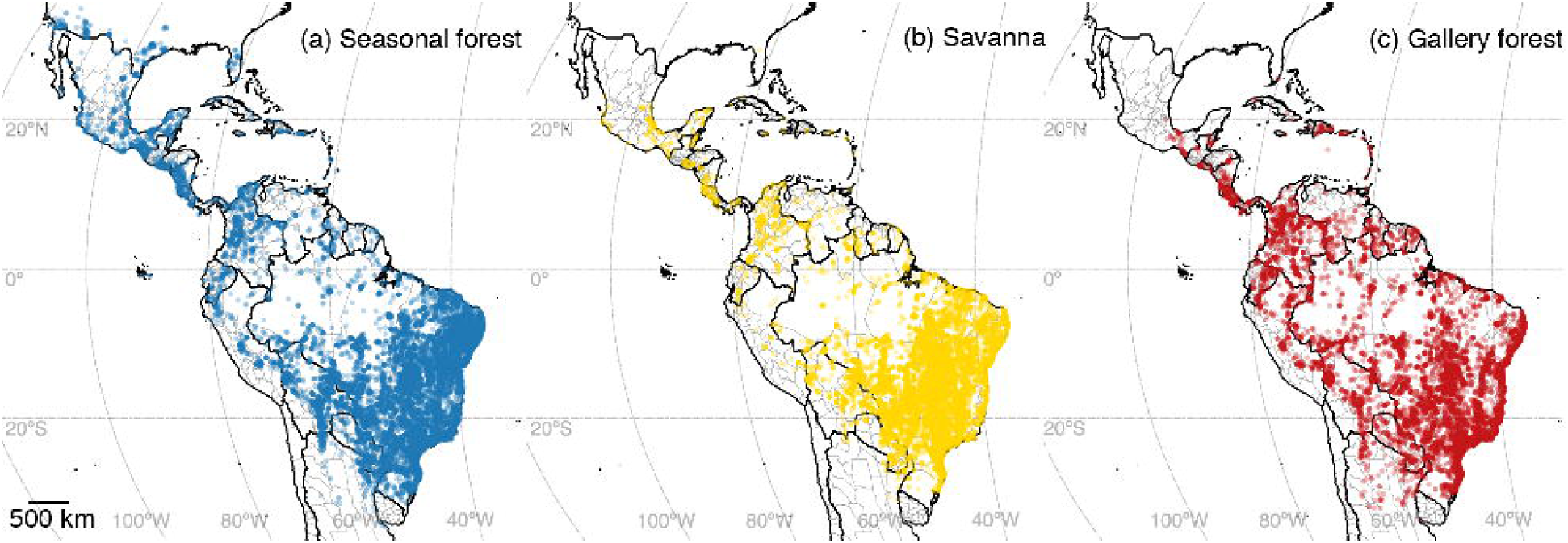
Geographic distribution points for 311 woody plants species associated with the three vegetation types included in the endemism analysis. (a) Distribution points of species of seasonally dry tropical forests; (b) distribution points of species of the savanna; and (c) distribution points of species of gallery forests.

We generated a map of weighted endemism using the algorithm proposed by Guerin, Ruokolainen, and Lowe (2015), applying a geographic grid with 0.25 × 0.25 degrees cells to generate the presence/absence matrix. We calculated the number of unique species that occurred in each cell and weighted this parameter for each species by dividing it by the number of grid cells in which they occurred (Guerin et al., 2015). By doing so, we expected to reduce the bias of widespread generalist species in the estimates of endemism.

### Models of past distribution of species and identification of stability nuclei

To evaluate the relative climatic stability between plateaus and valleys, we divided the entire Cerrado region into highlands (≥ 500 m elevation) and depressions (< 500 m elevation). We sampled 1000 random points from all 19 Worldclim bioclimatic variables (Hijmans, Cameron, Parra, Jones, & Jarvis, 2005) to calculate Pearson’s pairwise correlations and discarded highly correlated variables (≥ 0.75). The set of a11 non-correlated variables included the following: annual precipitation, precipitation seasonality, precipitation of the driest quarter, precipitation of the warmest quarter, precipitation of the coldest quarter, mean diurnal range, isothermality, temperature seasonality, maximum temperature of the warmest month, mean temperature of the wettest quarter, and mean temperature of the driest quarter. For past scenarios (mid-Holocene, Last Glacial Maximum, and Last Interglacial), we used variables from the Model for Interdisciplinary Research on Climate earth system models (MIROC-ESM) and the Community Climate System Model (CCSM4-scaled) climatic models (Hijmans et al., 2005). We randomly sampled 1000 points for each landscape unit and extracted values of 11 bioclimatic variables to compare the present environmental conditions between areas. To compare the climatic stability of plateaus and depressions, we sampled the same set of random points for past climatic scenarios (mid-Holocene, Last Glacial Maximum, and Last Interglacial; Hijmans et al., 2005). We then calculated the variances of each environmental variable of plateaus and valleys and compared them using a bootstrap approach by sampling 1000 repetitions of the variances and comparing the significance of outcomes with Welch’s t-test (Welch, 1938) for each variable in each landscape unit.

To test the stable/unstable climate hypothesis, we assessed which locations presented concentrated areas of stability of Cerrado vegetation during climatic cycles of the Pleistocene. We generated niche models to estimate the past distribution and stability nuclei of 311 species associated with gallery forests, seasonal forests, and savanna. We added the elevation layer to the set of 11 non-correlated environmental layers used in the variance calculation step. For past climatic scenarios, we applied the present elevation layer when projecting models to the Last Interglacial (LIG), Last Glacial Maximum (LGM), and mid-Holocene, assuming that terrain elevation did not change substantially along the modeled time series, because the final uplift of the Brazilian Central plateau dates from the late Miocene (da Silva, 1997). We averaged models between both scenarios (MIROC-SM and CCSM4) and presented a consensus model for each of these time periods. For the LIG (ca. 120,000 – 140,000 years BP), we adopted a projection of bioclimatic values from WorldClim following Otto-Bliesner et al. (2006).

We used the maximum entropy machine-learning algorithm Maxent v. 3.3.3 (Phillips, Anderson, & Schapire, 2006) to model the potential distribution for each species and for the simulated datasets. We processed models and bioclimatic layers using the R packages “dismo” (Hijmans, Phillips, Leathwick, & Elith, 2013) and “raster” (Hijmans & van Etten, 2014). For species distribution models, we used 1000 background points for each species to estimate background information with pseudo-absences. We folded the occurrence data for each species into 20% test and 80% training datasets and evaluated the model performance with AUC-ROC values. We converted present models into binary models by establishing a threshold based on the lower presence training (LPT) value, from which a presence point is recorded in the actual data for the present model (Pearson, Raxworthy, Nakamura, & Townsend Peterson, 2007). Then, we applied this same procedure to past scenarios using the present model threshold. We summarized information from the models by counting presence cells per species for each scenario reconstruction. To facilitate comparisons between species, we converted the values into percentages by dividing areas by the larger range registered for each species. This approach enabled the detection of the most likely climatic scenario in which a species reached its maximum distribution range as well as comparisons between its glacial and interglacial distribution. We also combined all binary models for each species and for the whole dataset to identify individual- and community-level stable areas.

### Endemism correlation with elevation, climatic stability, marginality, and soil fertility

To test the correlation of endemism with elevation and marginality, we extracted values of elevation and the distance from the margin of the Cerrado ecoregion shape (Olson *et al*., 2008) for 1000 random points. To estimate soil fertility, we applied the same approach used by Lehmann, Archibald, Hoffmann, and Bond (2011), in which a proxy is used as the measure of total exchangeable bases (cation exchange capacity of soil at 30 cm depth) from the SoilGrids database (Hengl *et al*. 2017). We extracted the values of endemism for these same points and fitted them into linear models using Pearson’s product-moment correlation test to estimate the significance of these associations. We then compared the p-values of each predictor to discuss their potential importance.

## RESULTS

### Species list and endemism calculation

We obtained 63,682 registries for 311 species, including all physiognomies of the Cerrado (Fig. 1). For individual species information see Appendix Tables S1.1, S1.2, and S1.3. Our estimates of weighted endemism for the Cerrado returned higher levels of endemism in seven Brazilian states: Minas Gerais, Goiás, Mato Grosso, Mato Grosso do Sul, São Paulo, Paraná, and Bahia. We identified 28 endemism nuclei (i.e., areas corresponding to 75% of the weighted endemism), namely: (1) Chapada dos Guimarães; (2) Serra do Caiapó/Taquari-Itiquira depression; (3) Serra de Maracajú; (4) Serra da Bodoquena; (5) Morrarias do Urucum; (6) Serrania San Luis; (7) Araguaia valley; (8) Serra do Roncador; (9) Verde - Itarumã rivers; (10) Três Lagoas; (11) Bataguassu; (12) Upper Paraná basin; (13) Serra das Divisões; (14) Serra de Botucatu/Serra da Canastra; (15) Serra Geral - Tibagi basin; (16) Southern Planalto Central; (17) Chapada dos Pilões; (18) Central Serra da Barcaça; (19) Serra Dourada; (20) Serra dos Pirineus; (21) Northern Bahia western plateau; (22) Chapada dos Veadeiros; (23) Serra do Espinhaço; (24) Serra do Cipó; (25) Chapada das Mangabeiras; (26) Serra do Estrondo; (27) Central Bahia western plateau; and (28) Southern Serra Geral de Goiás (Figs. 2a and 2b).

**Figure 2.**
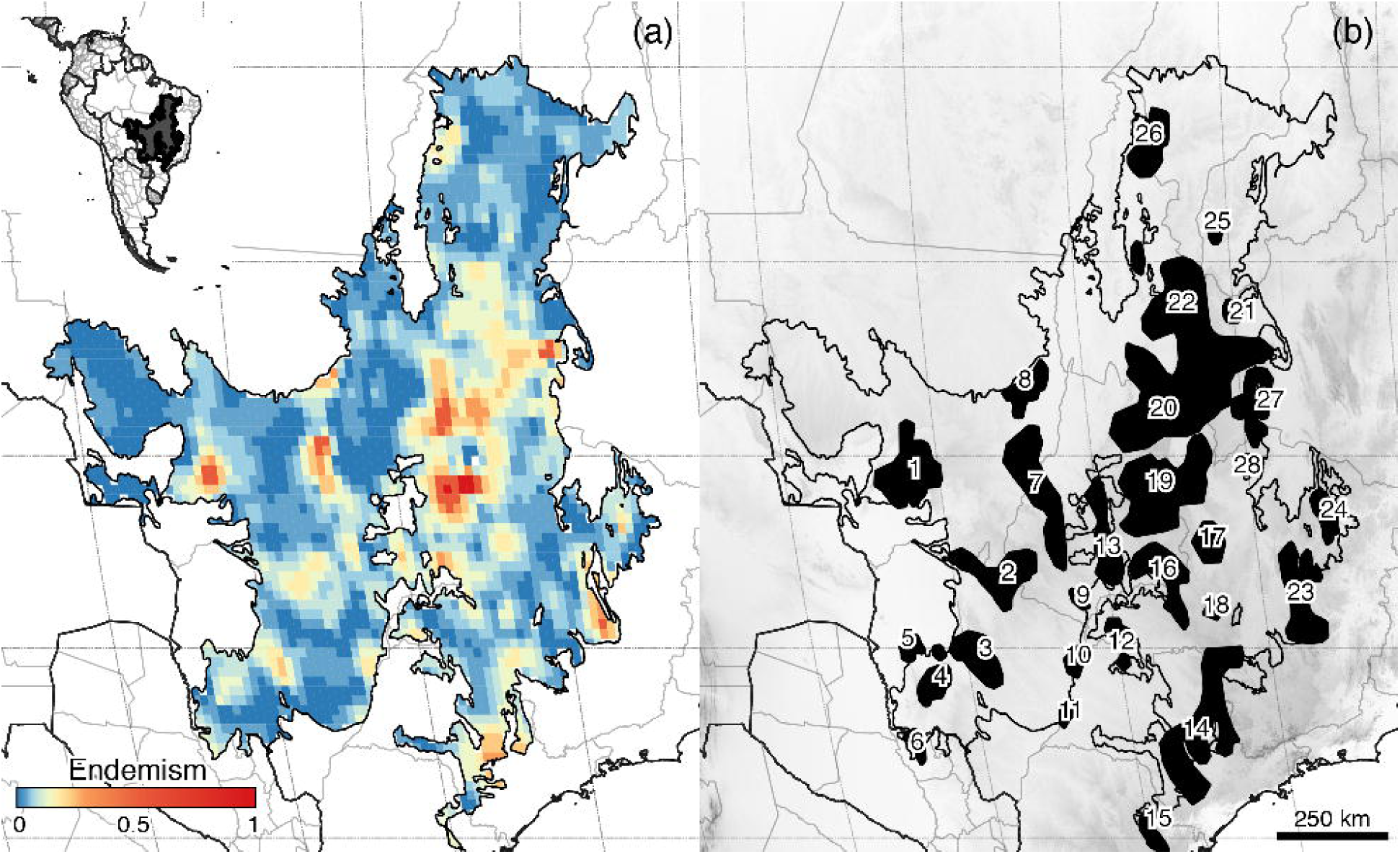
Distribution of 311 species of woody plants endemic to the Cerrado. (a) Weighted endemism (Guerín et al., 2015) map for woody species of the Cerrado domain. Grid cells are 0.25 × 0.25 degrees wide. (b) Corresponding areas containing the higher concentration (> 75%) of endemism for the Cerrado region: (1) Chapada dos Guimarães; (2) Serra do Caiapó/ Taquari-Itiquira Depression; (3) Serra de Maracajú; (4) Serra da Bodoquena; (5) Urucum Hills; (6) Serranía San Luis; (7) Araguaia Valley; (8) Serra do Roncador; (9) Verde - Itarumã Rivers; (10) Três Lagoas; (11) Bataguassu; (12) Upper Paraná Basin; (13) Serra das Divisões; (14) Serra de Botucatu/Serra da Canastra; (15) Serra Geral - Tibagi Basin; (16) Southern Planalto Central; (17) Chapada dos Pilões; (18) Serra da Barcaça; (19) Serra Dourada/Planalto Central; (20) Serra dos Pirineus; (21) Northern Chapada Ocidental da Bahia; (22) Chapada dos Veadeiros; (23) Serra do Espinhaço; (24) Serra do Cipó; (25) Chapada das Mangabeiras; (26) Serra do Estrondo; (27) Chapada Ocidental da Bahia; and (28) Southern Serra Geral de Goiás.

### Models of past distribution of species and identification of stability nuclei

The area under the curve for the receiver operating characteristics (AUC) for simulated datasets models was 0.736 for plateaus and 0.722 for valleys. For these data, our results recovered a pattern of glacial expansion for plateau environmental conditions, while valleys presented glacial retraction in terms of suitability (Figs. 3 and 4). In fact, the lowest proportional distribution for valleys and the highest distribution for plateaus occurred during the LGM (Fig. 4). For real occurrence data, species presented variable patterns (Supplementary Material – Tables S1.1, S1.2, and S1.3). A map of the distribution of multiple species stability nuclei was also obtained by combining models for stability regions of all species presence-absence maps along all climatic scenarios (Fig. 5).

**Figure 3.**
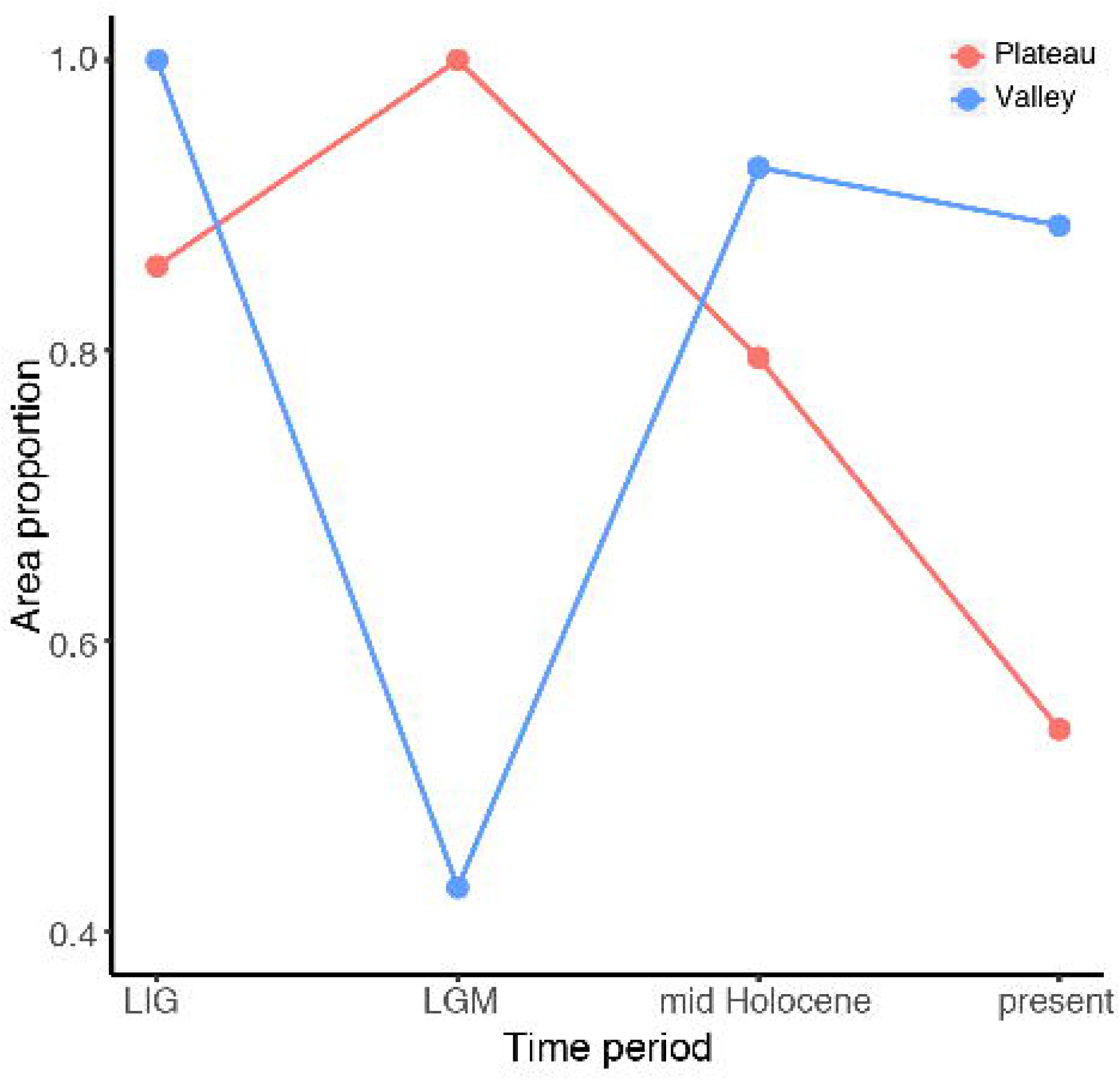
Potential environmental spaces for valley and plateau biotas based on simulated dataset models of presence/absence generated with Maxent (Phillips et al., 2006).

**Figure 4.**
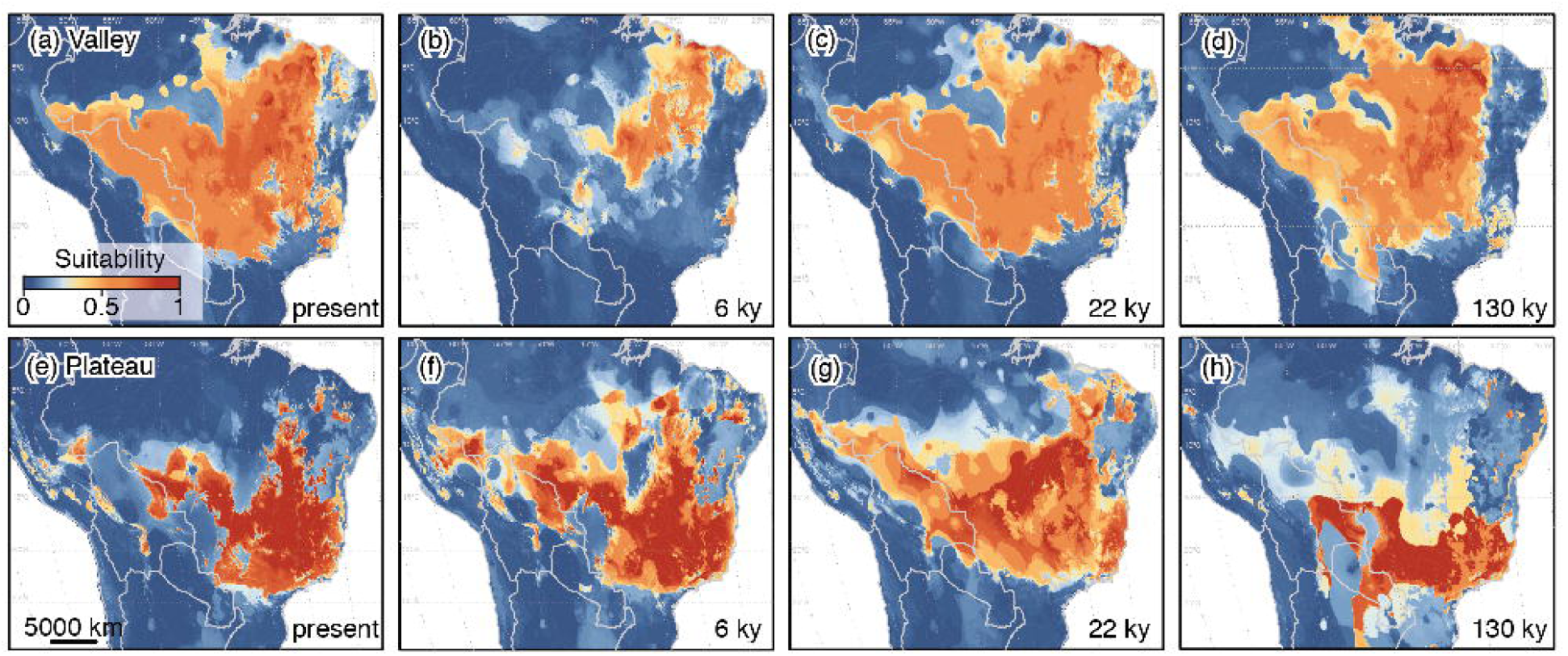
Potential suitable area for valley and plateau biotas of the Cerrado domain based on simulated dataset models of presence/absence generated with Maxent (Phillips et al., 2006). The area proportion is the relative size of the predicted presence compared to the maximum observed range for each landscape unit among the four modeled scenarios.

**Figure 5.**
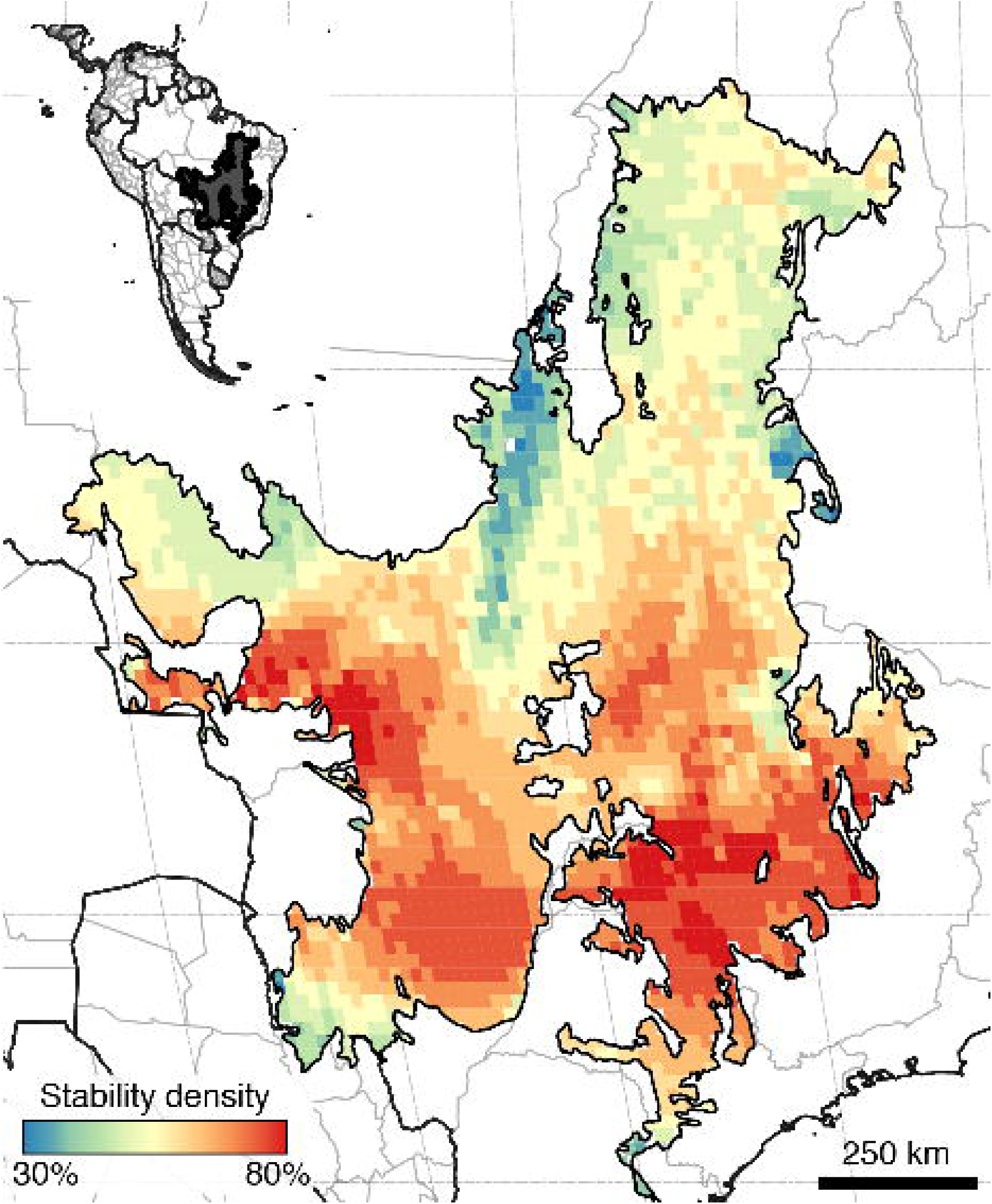
Density of range stability regions for 311 species of plants of the Cerrado region along four climatic scenarios for the past 130,000 years (present, mid Holocene, Last Glacial Maximum, and Last Interglacial).

### Endemism correlation with elevation, climatic stability, marginality of distribution, and soil fertility

We observed significant correlations (p-value < 0.01) for elevation and niche stability density; niche stability density and endemism; distance to the core and endemism; and elevation and endemism (Table 1; Figs. 6a, 6b, 6c, 6d). Linear models detected no correlation between soil fertility and endemism (p value > 0.01). Additionally, the generalized linear model, including all predictors, indicated that only elevation and marginality presented significant correlation with endemism levels (Table 1).

**Table 1.** Generalized linear model for endemism, elevation, marginality, and niche stability density among the Last Interglacial, Last Glacial Maximum, mid-Holocene and present (* p value < 0.01).

| Predictor | Estimate | Std. Error | t value | Pr(>\|t\|) | Significance |
| --- | --- | --- | --- | --- | --- |
| (Intercept) | $3.80 \times 10^{-2}$ | $1.87 \times 10^{-2}$ | 2.034 | 0.0422 | |
| Elevation | $9.20 \times 10^{-5}$ | $1.18 \times 10^{-5}$ | 7.8 | $\approx 0$ | * |
| Marginality | $-2.56 \times 10^{-7}$ | $4.16 \times 10^{-8}$ | -6.155 | $\approx 0$ | * |
| Soil Fertility | $2.28 \times 10^{-3}$ | $1.06 \times 10^{-3}$ | 2.143 | 0.0323 | |
| Stability density | $1.40 \times 10^{-2}$ | $2.58 \times 10^{-2}$ | 0.54 | 0.5891 | |

**Figure 6.**
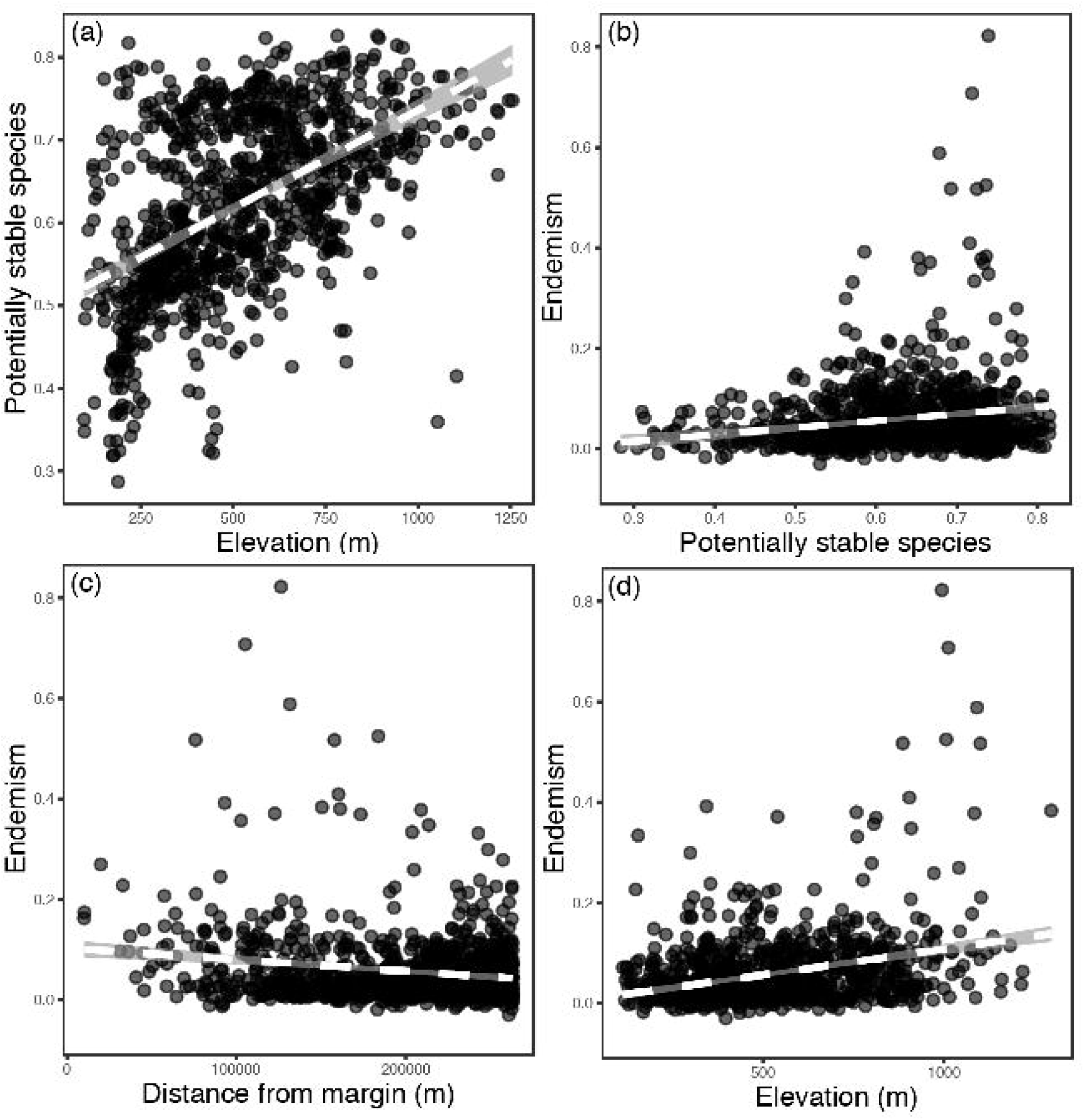
Linear models showing the correlation between weighted endemism, elevation, and distance to the core. (a) Stability density vs. elevation; (b) Endemism vs. Stability density; (c) Weighted endemism vs. distance to the margin; and (d) Weighted endemism vs. elevation.

All 11 non-collinear bioclimatic variables analyzed in this work presented significant differences between valleys and plateaus (Table 2). A total of 10 variables (ca. 90%) presented lower variances for plateaus than for valleys, suggesting that climatic conditions on plateaus were more stable during the investigated time periods (LIG, LGM, mid-Holocene). A single variable, precipitation seasonality, displayed lower variances in valleys. Welch’s two-sample t-test returned p-values lower than 0.01 for all comparisons.

**Table 2.** Mean values and confidence intervals of the variances of 11 bioclimatic variables for valley and plateau environments in the Brazilian Cerrado based on 1000 bootstrap replicates. Variances were calculated by comparing cell values for 1000 random points within each landscape unity for the present, mid-Holocene (6,000 years before present), Last Glacial Maximum (22,000 years before present), and Last Interglacial (130,000 years before present). *t-values*: t-values calculated with a paired t-test. (*) p-value > 0.01.

|  |  | Valleys |  | Plateaus |  |  |
| --- | --- | --- | --- | --- | --- | --- |
|  |  |  | 95% |  | 95% |  |
| Variable | Description | Mean | CI | Mean | CI | t-value |
| bio2 | Mean Diurnal Range | 132 | -19 | 113 | -20 | -221* |
| bio3 | Isothermality | 22 | -5 | 16 | -5 | -440* |
| bio4 | Temperature Seasonality | 285582 | -82242 | 203040 | -82843 | -538* |
| bio5 | Max. Temperature of Warmest Month | 569 | 47 | 615 | 46 | 152* |
| bio8 | Mean Temperature of Wettest Quarter | 282 | 47 | 329 | 46 | 287* |
| bio9 | Mean Temperature of Driest Quarter | 820 | -115 | 705 | -116 | -252* |
| bio12 | Annual Precipitation | 89629 | -18262 | 71271 | -18455 | -373* |
| bio15 | Precipitation Seasonality | 194 | -45 | 149 | -45 | -304* |
| bio17 | Precipitation of Driest Quarter | 1871 | -623 | 1245 | -630 | -338* |
| bio18 | Precipitation of Warmest Quarter | 46822 | -9568 | 37211 | -9655 | -431* |
| bio19 | Precipitation of Coldest Quarter | 58886 | -28905 | 29912 | -29043 | -821* |

## DISCUSSION

We generated endemism maps for Cerrado woody plants and compared the patterns observed with biogeographical theories of the Cerrado, which assumed plateau/depression landscape, climate stability/instability, and core/peripheral distribution, and low/high soil nutrient availability as central processes for the generation of endemism. We demonstrated that the climate presented an overall higher stability in the plateaus than in the valleys. A positive correlation between multiple-species stable suitability regions and endemism was observed. The soil fertility, however, showed weak correlation with endemism. Thus, we managed to link the distribution of higher endemism nuclei to niche stability density, elevation and centrality, reinforcing the importance of these environmental elements in the biogeographic processes that generated the Cerrado plants diversity. We then discuss our results in the context of climate changes, tectonic history, and biogeographic processes of the Cerrado.

### Elevation and marginality of distribution as predictors of endemism

We identified 28 areas harboring more than 75% of the endemism of the Cerrado flora. The identification of nuclei of endemism in some regions, such as Chapada dos Guimarães, Chapada dos Veadeiros, Serra do Cipó, and Planalto Central is consistent with endemism patterns identified for other taxa (Simon & Proença, 2000). Most of the identified endemism areas are located in regions in high elevation, whereas fewer valley regions, such as the Araguaia Valley, Itarumã, Três Lagoas, Upper Paraná Basin, Taquari-Itiquira Depression, Tibagi Basin, identified as highly endemic, supporting results of our correlation analysis. **Elevation** along with landscape compartmentalization are considered important drivers of diversification worldwide, as differences in elevation often promote isolation, a key process for speciation and endemism (MacArthur & Wilson, 1967; Chen & He, 2009; Slaton, 2015). Although the concentration of endemism levels of plants at high elevations was already demonstrated on a global scale (Steinbauer et al., 2016), we present here a finer scale pattern of endemism in the Cerrado, reinforcing the previously suggested relevance of this correlation in this specific ecosystem (e.g., Fiaschi & Pirani, 2009). Given the mosaic-like aspect of the topography of the Cerrado, mostly composed by plateaus separated from each other by interconnected depressions and river valleys, the higher concentration of endemism regions in altitudes can be interpreted as a result of the geographic isolation between the plateaus (Alves & Kolbek, 1994). Following the diversification model proposed by Chen and He (2009) and the classic island biogeography model by MacArthur and Wilson (1967), we propose that the degree of highlands connectivity over the Center-West region of Brazil, along with their geographic extent influence the proportion of the flora, which is shared between plateaus, and also the susceptibility of populations to stochastic local extinction events, directly influencing endemism levels of the plateaus. Under these assumptions, the geographic composition of the Cerrado constitutes a system that is similar to a mosaic of habitats in highlands, thus promoting the isolation of lineages in hilltops, which explains the higher concentration of endemic species in these landscapes.

In addition to elevation, the centrality of the distribution was also identified as an important predictor of endemism. This pattern can be explained by the fact that species that are adapted to a particular environment tend to present higher chances of survival, reproductive success and population growth under their optimal conditions (Eckert et al., 2008). These conditions are gradually distributed over the geographical space and are expected to be more suitable in nuclear areas of distributions. Although transition zones may occasionally show high species richness, this phenomenon is often caused by the presence of widespread species whose distributions overlap at ecotones (e.g. Brooks et al., 2001).

However, in certain cases, marginal environments could be subjected to contrasting environmental filters compared to adjacent core environments, such as the pattern reported in the Atlantic Rainforest, for which the geographical distribution of the endemic plants accounted for almost half of the total endemism of plants in the biome (Neves et al., 2017).

To the best of our knowledge, this is the first study that demonstrates this correlation for the Cerrado. Based on the results presented in this work, we suggest that marginal environments of the Cerrado are mostly occupied by widespread galleries of seasonal forest species, which are also commonly present in other tropical biomes, such as the Amazon and the Atlantic rainforests. Comparatively, core regions of the Cerrado harbor species with more restricted niche and distributions, often showing some level of habitat specialization (e.g., thick barks, deep roots systems, and sclerophylly) and mostly exclusively associated with savanna environments. The observed correlation between endemism and centrality of distribution may also be affected by the interaction of centrality with two other predictors tested in this work: the historical climatic suitability of nuclear regions for plants of the Cerrado (Bueno et al., 2017; Costa et al., 2018, Buzatti et al., 2017; Buzatti et al., 2018; Collevatti et al., 2018; Correa Ribeiro et al., 2016; Lima et al., 2017); and the concentration of higher elevation regions in core regions (e.g., Werneck, 2011).

### Endemism and niche stability

The positive correlation between **distribution stability** and **levels of endemism** is an indicator of the important role of environmental stability in maintaining the demographic stability of paleoendemic species of the Cerrado. The importance of regions of historical environmental stability for the assemblage and diversification of communities was already reported for the Cerrado (Werneck, 2011) and for other Neotropical biomes, such as the Amazon (Haffer, 1969, Bonaccorso *et al*., 2006) and the Atlantic rainforests (Carnaval & Moritz, 2008). According to Jansson (2003), the high levels of endemism observed in these stable nuclei is attributed to the fact that the lesser variable climatic conditions occurring in these environments favors both the persistence of paleoendemic species and the maintenance of sufficiently diverse gene pools, which allow for the diversification of neoendemics. Other lines of evidence based on paleo palynological and phylogeographic studies also support the relevance of historical stability of the nuclear regions of the Cerrado to species endemism. The uniform presence of charcoal particles deposition in the pollen records in core regions is a strong evidence of more stable presence of grass vegetation compared with marginal areas (Salgado-Labouriau, 1997; Pessenda et al. 1998). Phylogeographic studies show many examples of taxa in which core areas of the Cerrado present both higher demographic stability along the Pleistocene and higher genetic diversity in their populations (Correa Ribeiro et al., 2016; Buzatti et al., 2017; Lima et al. 2017; Collevatti et al., 2018). Such evidence may explain the increased environmental stability and endemism of nuclear regions of the Cerrado.

We also observed a positive correlation between the **density of stable species distribution** over the Quaternary and **elevation**. This pattern was also demonstrated by the result of the bootstrap analysis of stability, which recovered a higher climatic stability for most of the 11 variables assessed in highlands. Despite the relative stability of plateau environmental sets, we also demonstrated via simulated dataset models that **the responses of the climatic suitability of valley and plateau** points throughout the climatic shifts of the Quaternary presented opposite patterns of retraction and expansion. This result indicates that species associated with these environments are expected to have experienced distinct range shifts in response to these climatic fluctuations, as hypothesized by Ab’Saber (1982) and Werneck et al. (2011). These authors suggested that the ancient nuclear plateaus from Central Brazil, such as Chapada dos Guimarães and the Goiás Plateau, are likely to have constituted climatic refugia for several species of plants of the Cerrado during glacial and interglacial vegetation shifts and these processes may be responsible for the higher endemism levels observed in these areas.

### Role of soil characteristics and fire regime in shaping the distribution and diversification of the Cerrado flora

Among the environmental factors potentially shaping the distribution of plants endemism nuclei in the Cerrado, the role of edaphic factors and fire regimes must be considered because their effects have been demonstrated for several plants of this biome (Coutinho, 1990; Montgomery, 1988; Lehmann et al., 2014). Soil fertility and acidity are listed as potential factors influencing the patterns of distribution of species in the Cerrado (Ruggiero et al., 2002; Amorim & Batalha, 2006, 2007; da Silva & Batalha, 2008; Reatto et al., 2008; Dantas & Batalha, 2011; Lehmann et al., 2011; Staver et al., 2011; Franco et al., 2014). Using cation exchange as a potential predictor of soil fertility, we found no correlation between endemism and fertility levels, which means that the level of endemism in the Cerrado is undistinguishable between nutrient poor and rich soils. We suggest a trade-off of between endemic species diversification and extinction at both conditions of soil fertility. On the one hand, higher soil fertilities allow for the maintenance of higher population sizes and optimal chances of survival and reproduction; while on the other hand, poor nutrient soils represent strong abiotic filters and less intense competition among species, thus requiring the evolution of local adaptation. In this context, we speculate that endemics appear in both conditions: in the first case, they appear due to the stability of optimal conditions and, in the latter case, they appear due to the need of an adaptive response to environmental harshness. Still, both our results and interpretations are limited because soil conditions can be highly variable at finer scales, and important elements, such as specific nutrient availability or biologic interactions, may be absent from global interpolated datasets (Ettema & Wardle, 2002). We recommend the development of finer-scale studies to evaluate local effects of soil properties on plant communities composition in the Cerrado (e.g., Ruggiero et al., 2002; Amorim & Batalha, 2006; Amorim & Batalha, 2007), because such studies may assist in elucidating the role played by this environmental filter.

The historic occurrence of fire is also considered a key factor influencing the distribution of many species of plants in the Cerrado (Booysen & Tainton, 1984; Ruggiero et al. 2002; Amorim & Batalha, 2006; Amorim & Batalha, 2007; Staver et al., 2011; Lehmann et al., 2011, 2014) and the diversification of the endemic plant species in the Cerrado (Simon & Pennington, 2012). In a study evaluating the impacts of fire frequency in the diversity of plants in the Cerrado, da Silva and Batalha (2008) reported that although fire might reduce species richness in frequently burned sites, it drives the selection of fire-resistant species and is preferentially associated with these localities, and it excludes competitors. Thus, the historical recurrence of fire may have determined the persistence and distribution of fire-adapted endemic lineages during the biogeographic histories of various taxa, thereby resulting in higher endemism levels in frequently burned sites (Simon & Pennington, 2012; Neves et al., 2017). However, due to the lack of datasets describing the historical fire frequency for the Cerrado, testing this hypothesis may constitute a daunting challenge.

Possible approaches include the use of a fire-occurrence proxy, such as grass coverage percentual (ex. Neves et al., 2017), or even analyses of profiles of charcoal deposition in the pollen record descriptions available for the region (Pessenda *et al*., 1996; Salgado-Labouriau, 1997). Indeed, the uniform presence of charcoal particles in the palynological record for some sites suggests that the vegetation in core regions of the Cerrado, which harbors most of the endemic plants, presented recurrent occurrences of fire over at least 32,000 years before present (Salgado-Labouriau, 1997). Comparatively, some records from marginal regions, such as the Rondonia site, show more recent deposition of charcoal between 7,000 and 6,000 years before present (Pessenda *et al*. 1998), suggesting that the fire regime in these areas was probably established during the mid-Holocene. This suggests that the historical fire susceptibility of the core region of the Cerrado could also have enhanced the proportion of endemics species.

## CONCLUSION

We conclude that among the hypotheses discussed, elevation, stability, and centrality are key predictors of the distribution of endemic plants in the Cerrado. We identified that the majority of endemic plant species were concentrated in environmentally stable nuclear highland regions, whereas unstable and peripheral depressions present lower endemism levels. We interpreted these results as a product of a complex interaction between the diversification of neoendemic species via allopatric speciation in the plateaus and the persistence of paleoendemic lineages in climatically-stable nuclear areas. Our results are consistent with patterns reported for other plants in the literature, both for endemism and for intraspecific genetic diversity. Concerning soil fertility, our results suggest that the cation exchange capacity does not correlate with endemism levels, which are evenly distributed over high and low fertility soils. We also discussed the potential role of fire, which was not tested in our analyses but is recognized as an important environmental driver of patterns of species richness and endemism distributions in the Cerrado flora. The higher frequency of fire suggested in the literature based on palynological record in nuclear areas of the Cerrado may indicate the role of this environmental filter in endemism. However, both the soil and fire-frequency hypotheses require finer-scale tests for a deeper understanding of their relative importance.

Our results provide an overview of the geographic distribution of endemism of plants in a biogeographically complex system and may assist in directing conservation efforts and defining priority areas for conservation. Based on our results, we recommend that nuclear plateaus should be priority regions when delimiting protected areas for optimizing the inclusion of endemics species in the Cerrado. We recommend further studies on community-level approaches, as we implemented here, with other organisms, to test the extent of our interpretations.

## Supporting information

## Acknowledgements

This work was supported by the Coordenação de Aperfeiçoamento de Pessoal de Nível Superior (CAPES) in the form of a scholarship to J.D. Vidal. This research was also supported by CAPES - Computational Biology Program.

We would like to thank professors Fábio Pinheiro and Paulo De Marco Júnior for reading and suggesting important improvements to the manuscript.

## Author contributions

JDV designed the study, performed the analysis, and wrote the manuscript; and APS and IK assisted with the discussion of results and wrote the manuscript.

## Data accessibility statement

The weighted endemism raster data generated in this study are available as raster grids on the Pangaea database: https://issues.pangaea.de/browse/PDI-18934. The GBIF material list used for the endemism maps and models is available as supplementary material.

## Supporting Information

**Appendix S1.1**: List of modeled species for the seasonal forest.

**Appendix S1.2**: List of modeled species for the gallery forest.

**Appendix S1.3**: List of modeled species for the savanna.

**Appendix S2**: Sample points coordinates and GBIF identification used for endemism analysis and distribution models.

