## Supplementary material for "Impacts of landscape composition, marginality, and climatic stability on the patterns of endemism of Cerrado woody plants"

Supporting information

**Appendix S1.1**: List of modeled species for the seasonal forest.

| Species | N | AUC | Present | MH | LGM | LIG |
| --- | --- | --- | --- | --- | --- | --- |
| *Albizia inundata* | 19 | 0.9847 | 0.74 | 0.51 | 0.06 | 1 |
| *Alibertia edulis* | 371 | 0.8099 | 0.98 | 1 | 0.96 | 0.99 |
| *Alseis floribunda* | 241 | 0.8893 | 0.78 | 0.93 | 1 | 0.98 |
| *Amburana cearensis* | 296 | 0.8594 | 0.71 | 0.84 | 0.75 | 1 |
| *Anadenanthera colubrina* | 406 | 0.8374 | 0.59 | 0.86 | 0.74 | 1 |
| *Anadenanthera peregrina* | 384 | 0.8122 | 0.95 | 0.97 | 0.95 | 1 |
| *Aspidosperma cuspa* | 138 | 0.895 | 0.92 | 0.94 | 0.58 | 1 |
| *Aspidosperma discolor* | 124 | 0.9048 | 0.71 | 0.98 | 1 | 1 |
| *Aspidosperma polyneuron* | 177 | 0.898 | 0.92 | 0.92 | 1 | 0.91 |
| *Aspidosperma pyrifolium* | 395 | 0.8553 | 0.72 | 0.83 | 0.57 | 1 |
| *Aspidosperma riedelii* | 22 | 0.962 | 0.65 | 0.77 | 0.98 | 1 |
| *Aspidosperma subincanum* | 264 | 0.8508 | 0.79 | 0.92 | 0.85 | 1 |
| *Astronium fraxinifolium* | 298 | 0.8265 | 0.7 | 0.9 | 0.8 | 1 |
| *Balfourodendron riedelianum* | 164 | 0.9145 | 0.67 | 0.78 | 1 | 0.8 |
| *Bauhinia brevipes* | 223 | 0.864 | 0.89 | 0.92 | 1 | 0.99 |
| *Bowdichia virgilioides* | 358 | 0.8385 | 0.77 | 0.93 | 0.86 | 1 |
| *Brosimum gaudichaudii* | 346 | 0.8185 | 0.87 | 0.92 | 0.9 | 1 |
| *Brunfelsia australis* | 71 | 0.9393 | 0.71 | 0.71 | 1 | 0.63 |
| *Brunfelsia uniflora* | 253 | 0.8598 | 0.7 | 0.92 | 0.89 | 1 |
| *Cabralea canjerana* | 379 | 0.8437 | 0.71 | 0.81 | 1 | 0.76 |
| *Callisthene fasciculata* | 319 | 0.8393 | 0.78 | 0.87 | 0.8 | 1 |
| *Calycophyllum multiflorum* | 125 | 0.9144 | 1 | 0.97 | 0.92 | 0.9 |
| *Cariniana estrellensis* | 338 | 0.8595 | 0.52 | 0.9 | 1 | 0.85 |
| *Cassia ferruginea* | 177 | 0.8978 | 0.65 | 0.88 | 0.86 | 1 |
| *Cavanillesia umbellata* | 34 | 0.9645 | 0.84 | 1 | 0.86 | 0.96 |
| *Cedrela fissilis* | 345 | 0.8159 | 0.8 | 0.91 | 0.87 | 1 |
| *Celtis pubescens* | 159 | 0.875 | 0.8 | 0.87 | 1 | 0.93 |
| *Centrolobium tomentosum* | 79 | 0.9367 | 0.8 | 0.97 | 1 | 0.96 |
| *Chloroleucon tenuiflorum* | 106 | 0.9168 | 0.91 | 0.97 | 1 | 0.82 |
| *Combretum duarteanum* | 182 | 0.8986 | 0.83 | 0.92 | 0.53 | 1 |
| *Combretum leprosum* | 361 | 0.8489 | 0.77 | 0.89 | 0.6 | 1 |
| *Commiphora leptophloeos* | 387 | 0.8524 | 0.71 | 0.89 | 0.63 | 1 |
| *Cordia alliodora* | 343 | 0.8493 | 0.99 | 0.95 | 0.9 | 1 |
| *Cordia glazioviana* | 11 | 0.9754 | 0.51 | 0.38 | 0.49 | 1 |
| *Cordia trichotoma* | 358 | 0.8343 | 0.68 | 0.92 | 0.84 | 1 |
| *Cordiera sessilis* | 115 | 0.8795 | 0.9 | 0.99 | 0.99 | 1 |
| *Couepia uiti* | 82 | 0.9265 | 0.63 | 0.9 | 0.2 | 1 |
| *Coutarea hexandra* | 345 | 0.8345 | 0.8 | 0.94 | 0.86 | 1 |
| *Diatenopteryx sorbifolia* | 117 | 0.9244 | 0.83 | 0.9 | 1 | 0.83 |
| *Dilodendron bipinnatum* | 164 | 0.8905 | 0.72 | 0.9 | 0.91 | 1 |
| *Diplokeleba floribunda* | 66 | 0.9549 | 0.62 | 1 | 0.44 | 0.81 |
| *Dipteryx alata* | 314 | 0.8385 | 0.91 | 0.93 | 0.88 | 1 |
| *Duguetia furfuracea* | 338 | 0.8551 | 0.56 | 0.8 | 0.9 | 1 |
| *Emmotum nitens* | 342 | 0.8393 | 0.95 | 0.95 | 0.96 | 1 |
| *Enterolobium contortisiliquum* | 352 | 0.834 | 0.93 | 0.95 | 0.9 | 1 |
| *Fraunhofera multiflora* | 56 | 0.9713 | 0.59 | 0.53 | 0.43 | 1 |
| *Guazuma ulmifolia* | 309 | 0.8567 | 0.96 | 1 | 0.82 | 0.98 |
| *Helicteres brevispira* | 22 | 0.9236 | 0.7 | 1 | 1 | 0.82 |
| *Hirtella glandulosa* | 307 | 0.8405 | 0.78 | 0.91 | 0.88 | 1 |
| *Hymenaea courbaril* | 406 | 0.812 | 0.9 | 0.92 | 0.9 | 1 |
| *Hymenaea eriogyne* | 110 | 0.9394 | 0.71 | 0.3 | 0.67 | 1 |
| *Ipomoea carnea* | 315 | 0.8567 | 0.91 | 0.94 | 0.72 | 1 |
| *Jacaranda brasiliana* | 183 | 0.891 | 0.78 | 0.89 | 0.82 | 1 |
| *Jacaranda caroba* | 190 | 0.9177 | 0.41 | 0.6 | 1 | 0.74 |
| *Lafoensia pacari* | 349 | 0.8436 | 0.65 | 0.86 | 0.84 | 1 |
| *Lithraea molleoides* | 313 | 0.8649 | 0.62 | 0.89 | 1 | 0.91 |
| *Lonchocarpus sericeus* | 68 | 0.9538 | 0.6 | 0.96 | 0.5 | 1 |
| *Luehea candicans* | 288 | 0.8222 | 0.81 | 0.9 | 0.88 | 1 |
| *Luehea paniculata* | 232 | 0.8459 | 0.82 | 0.92 | 0.89 | 1 |
| *Machaerium acutifolium* | 372 | 0.8312 | 0.81 | 0.91 | 0.88 | 1 |
| *Machaerium villosum* | 141 | 0.9135 | 0.7 | 0.63 | 1 | 0.56 |
| *Magonia pubescens* | 263 | 0.8568 | 0.85 | 0.91 | 0.88 | 1 |
| *Miconia albicans* | 355 | 0.8364 | 0.78 | 0.92 | 0.89 | 1 |
| *Miconia macrothyrsa* | 201 | 0.8607 | 0.98 | 0.96 | 1 | 0.93 |
| *Monteverdia truncata* | 53 | 0.9455 | 0.73 | 1 | 0.75 | 0.84 |
| *Muellera montana* | 30 | 0.9817 | 0.78 | 0.27 | 1 | 0.18 |
| *Myracrodruon urundeuva* | 405 | 0.9345 | 0.72 | 1 | 0.57 | 0.91 |
| *Myroxylon balsamum* | 242 | 0.8621 | 0.99 | 0.82 | 1 | 0.76 |
| *Parkinsonia aculeata* | 396 | 0.8372 | 0.91 | 0.94 | 0.91 | 1 |
| *Peltophorum dubium* | 338 | 0.9655 | 0.69 | 0.73 | 0.39 | 1 |
| *Phyllostylon rhamnoides* | 120 | 0.8384 | 0.71 | 0.95 | 0.84 | 1 |
| *Physocalymma scaberrimum* | 208 | 0.9291 | 0.96 | 1 | 0.51 | 0.76 |
| *Phytolacca dioica* | 214 | 0.858 | 1 | 0.82 | 0.85 | 0.75 |
| *Piptadenia viridiflora* | 291 | 0.89 | 0.61 | 0.8 | 1 | 0.86 |
| *Plathymenia reticulata* | 369 | 0.8627 | 0.92 | 0.95 | 0.91 | 1 |
| *Platycyamus regnellii* | 59 | 0.8366 | 0.83 | 0.89 | 0.85 | 1 |
| *Platypodium elegans* | 430 | 0.9697 | 0.61 | 0.7 | 1 | 0.4 |
| *Poeppigia procera* | 312 | 0.828 | 0.78 | 0.88 | 0.89 | 1 |
| *Pouteria gardneriana* | 112 | 0.8783 | 0.95 | 0.89 | 1 | 0.88 |
| *Pseudobombax tomentosum* | 67 | 0.8892 | 0.75 | 1 | 0.77 | 0.81 |
| *Psychotria hoffmannseggiana* | 330 | 0.9333 | 0.86 | 1 | 0.84 | 0.47 |
| *Pterogyne nitens* | 326 | 0.8524 | 0.75 | 0.92 | 0.79 | 1 |
| *Qualea grandiflora* | 361 | 0.8339 | 0.71 | 0.88 | 0.89 | 1 |
| *Rauvolfia ligustrina* | 159 | 0.9166 | 0.73 | 0.85 | 0.6 | 1 |
| *Rudgea viburnoides* | 310 | 0.8471 | 0.67 | 0.81 | 1 | 0.7 |
| *Ruprechtia laxiflora* | 259 | 0.8676 | 0.71 | 0.96 | 0.93 | 1 |
| *Schinopsis brasiliensis* | 355 | 0.8711 | 0.7 | 0.91 | 0.84 | 1 |
| *Senna spectabilis* | 360 | 0.8533 | 0.76 | 0.93 | 1 | 0.94 |
| *Sideroxylon obtusifolium* | 343 | 0.8678 | 0.83 | 0.96 | 0.83 | 1 |
| *Siphoneugena densiflora* | 114 | 0.9438 | 0.49 | 0.61 | 1 | 0.41 |
| *Solanum granulosoleprosum* | 288 | 0.8775 | 0.76 | 0.96 | 1 | 0.94 |
| *Spondias tuberosa* | 312 | 0.8784 | 0.59 | 0.88 | 0.62 | 1 |
| *Sterculia striata* | 191 | 0.8585 | 0.83 | 0.92 | 0.8 | 1 |
| *Tabebuia aurea* | 27 | 0.9442 | 0.67 | 1 | 0.53 | 0.84 |
| *Tachigali peruviana* | 323 | 0.8483 | 0.73 | 0.87 | 0.85 | 1 |
| *Trichilia elegans* | 383 | 0.8361 | 0.75 | 0.82 | 0.89 | 1 |
| *Varronia leucocephala* | 67 | 0.9535 | 0.48 | 0.51 | 0.61 | 1 |
| *Vasconcellea quercifolia* | 25 | 0.9382 | 0.64 | 0.83 | 1 | 0.87 |
| *Vochysia haenkeana* | 185 | 0.8607 | 0.86 | 0.91 | 0.93 | 1 |
| *Xylopia aromatica* | 339 | 0.8122 | 0.99 | 0.99 | 0.99 | 1 |
| *Zanthoxylum rhoifolium* | 398 | 0.8075 | 0.82 | 0.92 | 0.95 | 1 |
| *Ziziphus joazeiro* | 16 | 0.9583 | 1 | 0.61 | 0.1 | 0.68 |

**Appendix S1.2**: List of modeled species for the gallery forest.

| Species | N | AUC | Present | MH | LGM | LIG |
| --- | --- | --- | --- | --- | --- | --- |
| *Acosmium cardenasii* | 53 | 0.9578 | 1 | 0.99 | 0.83 | 0.76 |
| *Annona aurantiaca* | 48 | 0.964 | 1 | 0.69 | 0.19 | 0.6 |
| *Apeiba tibourbou* | 336 | 0.81 | 0.98 | 1 | 0.92 | 0.99 |
| *Apuleia leiocarpa* | 314 | 0.8257 | 0.89 | 0.96 | 0.91 | 1 |
| *Aspidosperma quebracho-blanco* | 126 | 0.9003 | 0.87 | 1 | 0.88 | 0.97 |
| *Bauhinia mollis* | 124 | 0.8946 | 0.77 | 1 | 0.81 | 0.86 |
| *Callisthene major* | 166 | 0.9164 | 0.89 | 0.97 | 1 | 0.99 |
| *Callisthene mollissima* | 31 | 0.9719 | 0.86 | 1 | 1 | 0.57 |
| *Calophyllum brasiliense* | 356 | 0.8128 | 0.93 | 0.98 | 0.94 | 1 |
| *Cardiopetalum calophyllum* | 172 | 0.8942 | 0.99 | 0.78 | 1 | 0.51 |
| *Cariniana rubra* | 80 | 0.9303 | 1 | 0.92 | 0.67 | 0.83 |
| *Cheiloclinium cognatum* | 309 | 0.8305 | 0.91 | 0.91 | 0.89 | 1 |
| *Copaifera langsdorffii* | 333 | 0.8328 | 0.62 | 0.9 | 0.83 | 1 |
| *Copaifera oblongifolia* | 79 | 0.9389 | 0.81 | 1 | 0.88 | 0.76 |
| *Croton urucurana* | 333 | 0.8331 | 0.91 | 0.99 | 0.98 | 1 |
| *Cupania vernalis* | 358 | 0.8367 | 0.67 | 0.91 | 0.84 | 1 |
| *Dendropanax cuneatus* | 306 | 0.8525 | 0.65 | 0.72 | 1 | 0.67 |
| *Erythroxylum daphnites* | 177 | 0.8794 | 0.78 | 0.93 | 0.97 | 1 |
| *Eugenia ternatifolia* | 36 | 0.9616 | 0.42 | 1 | 0.63 | 0.77 |
| *Euplassa inaequalis* | 106 | 0.9279 | 0.92 | 1 | 1 | 0.95 |
| *Euterpe edulis* | 173 | 0.9142 | 0.41 | 0.88 | 1 | 0.81 |
| *Ferdinandusa speciosa* | 138 | 0.9307 | 0.82 | 0.89 | 1 | 0.92 |
| *Guarea guidonia* | 278 | 0.8126 | 0.89 | 0.93 | 0.9 | 1 |
| *Guarea kunthiana* | 347 | 0.8239 | 0.91 | 0.98 | 1 | 1 |
| *Guarea macrophylla* | 360 | 0.8348 | 0.92 | 0.93 | 0.96 | 1 |
| *Guatteria sellowiana* | 204 | 0.9152 | 0.51 | 0.7 | 1 | 0.66 |
| *Guettarda viburnoides* | 340 | 0.8382 | 0.59 | 0.88 | 0.77 | 1 |
| *Hedyosmum brasiliense* | 324 | 0.887 | 0.41 | 0.7 | 1 | 0.69 |
| *Ladenbergia cujabensis* | 11 | 0.9682 | 1 | 0.83 | 0.6 | 0.52 |
| *Licania apetala* | 271 | 0.8 | 0.98 | 0.96 | 0.92 | 1 |
| *Luetzelburgia praecox* | 21 | 0.9748 | 0.76 | 1 | 0.1 | 0.36 |
| *Lycium cuneatum* | 30 | 0.958 | 0.88 | 0.81 | 0.67 | 1 |
| *Machaerium eriocarpum* | 26 | 0.9726 | 0.66 | 0.97 | 0.02 | 1 |
| *Magnolia ovata* | 125 | 0.9578 | 0.35 | 0.57 | 1 | 0.36 |
| *Matayba guianensis* | 349 | 0.8164 | 0.91 | 0.98 | 0.94 | 1 |
| *Mauritia flexuosa* | 65 | 0.8384 | 1 | 0.92 | 0.8 | 0.93 |
| *Miconia chartacea* | 171 | 0.9252 | 0.59 | 0.83 | 1 | 0.79 |
| *Mimosa glutinosa* | 25 | 0.9856 | 0 | 0 | 1 | 0 |
| *Myrcia camapuanensis* | 28 | 0.9708 | 0.76 | 1 | 0.46 | 0.31 |
| *Myrcia fenzliana* | 62 | 0.9545 | 0.51 | 0.71 | 1 | 0.71 |
| *Myrcia racemulosa* | 26 | 0.9729 | 0.6 | 1 | 0.99 | 0.62 |
| *Myrcia splendens* | 345 | 0.8243 | 0.87 | 0.92 | 0.91 | 1 |
| *Myrcia tomentosa* | 27 | 0.985 | 0.94 | 0.93 | 1 | 0.42 |
| *Ocotea aciphylla* | 279 | 0.8731 | 0.93 | 1 | 0.98 | 0.96 |
| *Ouratea castaneifolia* | 357 | 0.7981 | 0.88 | 0.92 | 0.89 | 1 |
| *Piptocarpha macropoda* | 162 | 0.9223 | 0.42 | 0.61 | 1 | 0.55 |
| *Posoqueria latifolia* | 339 | 0.8362 | 0.91 | 0.96 | 0.93 | 1 |
| *Protium altissimum* | 274 | 0.8442 | 0.94 | 0.94 | 0.91 | 1 |
| *Protium heptaphyllum* | 295 | 0.8275 | 0.89 | 0.93 | 0.89 | 1 |
| *Pseudobombax minimum* | 15 | 0.9799 | 1 | 0.95 | 0.97 | 0.7 |
| *Pseudolmedia laevigata* | 295 | 0.8447 | 1 | 1 | 0.96 | 1 |
| *Psychotria carthagenensis* | 378 | 0.8324 | 0.85 | 0.92 | 0.93 | 1 |
| *Pterodon emarginatus* | 113 | 0.9208 | 0.91 | 0.77 | 1 | 0.45 |
| *Richeria grandis* | 341 | 0.8276 | 0.92 | 0.92 | 0.92 | 1 |
| *Schefflera morototoni* | 353 | 0.7908 | 0.95 | 0.98 | 0.95 | 1 |
| *Senegalia praecox* | 134 | 0.9111 | 1 | 0.97 | 0.96 | 0.68 |
| *Styrax camporum* | 278 | 0.8669 | 0.65 | 0.87 | 1 | 0.91 |
| *Symplocos nitens* | 146 | 0.9128 | 0.5 | 0.87 | 0.98 | 1 |
| *Talisia subalbens* | 11 | 0.9723 | 0.75 | 0.8 | 1 | 0.68 |
| *Tapirira guianensis* | 232 | 0.8396 | 0.92 | 0.95 | 0.92 | 1 |
| *Tapura amazonica* | 277 | 0.8243 | 0.96 | 0.94 | 0.94 | 1 |
| *Tetragastris balsamifera* | 38 | 0.9566 | 0.71 | 0.92 | 0.69 | 1 |
| *Virola sebifera* | 343 | 0.8237 | 1 | 0.75 | 0.99 | 0.68 |
| *Virola urbaniana* | 19 | 0.9868 | 0.66 | 1 | 0.75 | 0.11 |
| *Vochysia pruinosa* | 44 | 0.9743 | 0.57 | 0.24 | 1 | 0.24 |
| *Vochysia pyramidalis* | 242 | 0.8901 | 0.82 | 0.93 | 0.89 | 1 |
| *Vochysia tucanorum* | 312 | 0.8763 | 0.69 | 0.85 | 1 | 0.94 |
| *Xylopia emarginata* | 142 | 0.8595 | 0.96 | 1 | 1 | 0.88 |
| *Xylopia sericea* | 258 | 0.8263 | 0.85 | 0.9 | 0.87 | 1 |

**Appendix S1.3**: List of modeled species for the savanna.

| Species | N | AUC | Present | MH | LGM | LIG |
| --- | --- | --- | --- | --- | --- | --- |
| *Acrocomia aculeata* | 297 | 0.8518 | 0.8 | 0.91 | 1 | 0.96 |
| *Aegiphila verticillata* | 272 | 0.8566 | 0.91 | 1 | 0.82 | 0.81 |
| *Agonandra brasiliensis* | 277 | 0.8695 | 0.61 | 0.85 | 0.92 | 1 |
| *Allagoptera campestris* | 267 | 0.8213 | 0.92 | 0.97 | 0.89 | 1 |
| *Allagoptera leucocalyx* | 107 | 0.885 | 0.6 | 0.9 | 0.86 | 1 |
| *Anacardium humile* | 287 | 0.9038 | 0.66 | 0.95 | 0.88 | 1 |
| *Anacardium occidentale* | 324 | 0.8691 | 0.79 | 0.93 | 0.92 | 1 |
| *Andira cordata* | 70 | 0.8229 | 0.92 | 0.91 | 0.86 | 1 |
| *Andira cujabensis* | 165 | 0.9519 | 0.74 | 0.52 | 0.66 | 1 |
| *Andira vermifuga* | 161 | 0.9005 | 0.95 | 1 | 0.91 | 0.9 |
| *Annona coriacea* | 341 | 0.8731 | 0.8 | 0.87 | 0.84 | 1 |
| *Annona crassiflora* | 184 | 0.8371 | 0.75 | 0.91 | 1 | 0.99 |
| *Annona monticola* | 99 | 0.8941 | 0.65 | 0.64 | 1 | 0.63 |
| *Annona nutans* | 118 | 0.9277 | 0.77 | 0.93 | 1 | 0.98 |
| *Annona tomentosa* | 160 | 0.9344 | 0.82 | 1 | 0.49 | 0.85 |
| *Aspidosperma macrocarpon* | 287 | 0.9142 | 0.81 | 0.96 | 1 | 0.99 |
| *Aspidosperma multiflorum* | 126 | 0.8301 | 0.89 | 0.85 | 1 | 0.9 |
| *Aspidosperma nobile* | 66 | 0.8759 | 0.72 | 0.86 | 0.91 | 1 |
| *Aspidosperma tomentosum* | 296 | 0.928 | 1 | 0.87 | 0.5 | 0.73 |
| *Attalea speciosa* | 19 | 0.8531 | 0.79 | 0.99 | 0.98 | 1 |
| *Bauhinia rufa* | 343 | 0.9202 | 0.7 | 0.83 | 0.64 | 1 |
| *Butia archeri* | 34 | 0.8442 | 0.73 | 0.87 | 0.93 | 1 |
| *Byrsonima basiloba* | 192 | 0.9816 | 0.3 | 0.36 | 1 | 0.22 |
| *Byrsonima coccolobifolia* | 346 | 0.9231 | 0.62 | 0.54 | 1 | 0.42 |
| *Byrsonima intermedia* | 344 | 0.8403 | 0.82 | 0.99 | 1 | 0.87 |
| *Byrsonima pachyphylla* | 67 | 0.8595 | 0.54 | 0.87 | 0.79 | 1 |
| *Byrsonima verbascifolia* | 382 | 0.947 | 0.81 | 0.83 | 1 | 0.66 |
| *Campomanesia adamantium* | 313 | 0.8333 | 0.74 | 1 | 0.94 | 0.99 |
| *Campomanesia pubescens* | 280 | 0.8728 | 0.83 | 0.95 | 0.97 | 1 |
| *Caryocar brasiliense* | 67 | 0.8887 | 0.52 | 0.81 | 1 | 0.65 |
| *Caryocar coriaceum* | 88 | 0.9123 | 0.66 | 0.8 | 1 | 0.8 |
| *Caryocar cuneatum* | 27 | 0.9372 | 0.59 | 0.59 | 0.56 | 1 |
| *Cecropia pachystachya* | 395 | 0.9741 | 0.5 | 0.53 | 0.37 | 1 |
| *Chamaecrista machaeriifolia* | 33 | 0.8166 | 0.75 | 0.9 | 0.86 | 1 |
| *Chamaecrista orbiculata* | 176 | 0.9832 | 0.26 | 0.25 | 1 | 0.21 |
| *Cochlospermum regium* | 293 | 0.9225 | 0.89 | 0.78 | 1 | 0.59 |
| *Connarus detersus* | 24 | 0.9269 | 0.75 | 0.87 | 0.82 | 1 |
| *Connarus suberosus* | 292 | 0.8225 | 0.66 | 0.89 | 0.88 | 1 |
| *Copaifera elliptica* | 28 | 0.9406 | 0.83 | 0.89 | 0.91 | 1 |
| *Copaifera malmei* | 28 | 0.8486 | 0.89 | 0.87 | 0.42 | 1 |
| *Cordia insignis* | 90 | 0.9569 | 1 | 0.73 | 0.13 | 0.31 |
| *Couepia grandiflora* | 121 | 0.9683 | 0.49 | 0.84 | 0.63 | 1 |
| *Curatella americana* | 359 | 0.9259 | 0.72 | 0.82 | 1 | 1 |
| *Dalbergia miscolobium* | 296 | 0.9094 | 0.85 | 0.99 | 0.69 | 1 |
| *Davilla elliptica* | 342 | 0.8233 | 0.66 | 0.85 | 0.9 | 1 |
| *Davilla grandiflora* | 134 | 0.8804 | 0.71 | 0.87 | 0.88 | 1 |
| *Dimorphandra gardneriana* | 224 | 0.8553 | 0.61 | 0.93 | 0.89 | 1 |
| *Dimorphandra mollis* | 289 | 0.9112 | 0.77 | 0.78 | 0.8 | 1 |
| *Diospyros lasiocalyx* | 310 | 0.8727 | 0.84 | 0.9 | 0.93 | 1 |
| *Diptychandra aurantiaca* | 190 | 0.8503 | 0.61 | 0.88 | 0.79 | 1 |
| *Enterolobium gummiferum* | 135 | 0.8383 | 0.81 | 0.99 | 0.98 | 1 |
| *Eremanthus capitatus* | 13 | 0.8393 | 0.53 | 0.65 | 1 | 0.58 |
| *Eremanthus glomerulatus* | 192 | 0.922 | 0.54 | 0.45 | 0.6 | 1 |
| *Eremanthus mattogrossensis* | 43 | 0.9416 | 0.76 | 0.91 | 1 | 0.6 |
| *Eriotheca gracilipes* | 208 | 0.9572 | 0.75 | 0.97 | 1 | 0.94 |
| *Erythroxylum cuneifolium* | 242 | 0.8672 | 0.63 | 0.91 | 0.98 | 1 |
| *Erythroxylum engleri* | 64 | 0.8655 | 0.88 | 0.92 | 1 | 0.36 |
| *Erythroxylum suberosum* | 347 | 0.9486 | 0.69 | 0.92 | 0.91 | 1 |
| *Erythroxylum tortuosum* | 136 | 0.8497 | 0.66 | 0.65 | 1 | 0.46 |
| *Eschweilera nana* | 154 | 0.9202 | 0.88 | 0.96 | 0.91 | 1 |
| *Esenbeckia pumila* | 136 | 0.908 | 0.77 | 0.74 | 1 | 0.56 |
| *Eugenia dysenterica* | 267 | 0.9428 | 0.63 | 0.84 | 0.78 | 1 |
| *Ferdinandusa elliptica* | 176 | 0.9323 | 1 | 0.99 | 0.98 | 0.94 |
| *Hancornia speciosa* | 336 | 0.8348 | 0.74 | 0.88 | 0.83 | 1 |
| *Handroanthus ochraceus* | 79 | 0.8418 | 0.78 | 0.77 | 1 | 0.47 |
| *Himatanthus obovatus* | 334 | 0.8366 | 0.78 | 0.9 | 0.62 | 1 |
| *Hirtella ciliata* | 302 | 0.8651 | 0.74 | 0.88 | 0.9 | 1 |
| *Hymenaea stigonocarpa* | 372 | 0.8266 | 0.86 | 0.95 | 0.99 | 1 |
| *Hyptis campestris* | 88 | 0.9136 | 0.78 | 0.93 | 0.91 | 1 |
| *Ilex affinis* | 237 | 0.8779 | 0.69 | 0.92 | 1 | 0.92 |
| *Jacaranda decurrens* | 59 | 0.9429 | 0.77 | 0.89 | 0.92 | 1 |
| *Kielmeyera coriacea* | 327 | 0.8554 | 0.43 | 0.48 | 1 | 0.3 |
| *Kielmeyera grandiflora* | 79 | 0.9581 | 0.59 | 0.78 | 1 | 0.43 |
| *Kielmeyera rosea* | 26 | 0.9582 | 0.73 | 0.85 | 1 | 0.95 |
| *Kielmeyera rubriflora* | 273 | 0.8743 | 0.66 | 0.41 | 1 | 0.15 |
| *Kielmeyera speciosa* | 66 | 0.9649 | 0.68 | 0.76 | 1 | 0.87 |
| *Leptolobium dasycarpum* | 467 | 0.9504 | 0.59 | 0.72 | 1 | 0.72 |
| *Licania dealbata* | 95 | 0.8459 | 0.7 | 0.98 | 0.63 | 1 |
| *Licania humilis* | 111 | 0.9143 | 0.53 | 0.62 | 1 | 0.27 |
| *Luetzelburgia auriculata* | 172 | 0.8291 | 0.64 | 0.7 | 1 | 0.49 |
| *Lychnophora ericoides* | 248 | 0.8568 | 0.48 | 0.88 | 1 | 0.79 |
| *Machaerium opacum* | 164 | 0.9377 | 0.89 | 0.95 | 0.44 | 1 |
| *Maprounea brasiliensis* | 89 | 0.8164 | 0.87 | 1 | 0.21 | 0.37 |
| *Martiodendron mediterraneum* | 87 | 0.9257 | 0.62 | 0.88 | 1 | 0.97 |
| *Mezilaurus crassiramea* | 34 | 0.8769 | 0.44 | 0.49 | 1 | 0.57 |
| *Miconia chamissois* | 294 | 0.8977 | 0.63 | 0.47 | 1 | 0.36 |
| *Miconia theizans* | 284 | 0.9188 | 0.32 | 0.18 | 1 | 0.1 |
| *Mimosa claussenii* | 231 | 0.9887 | 0.4 | 0.66 | 0.01 | 1 |
| *Mimosa decorticans* | 16 | 0.9875 | 0.67 | 1 | 0.82 | 0.99 |
| *Mimosa interrupta* | 20 | 0.9444 | 0.46 | 0.59 | 1 | 0.59 |
| *Mimosa laticifera* | 59 | 0.9675 | 0.8 | 1 | 0.94 | 0.62 |
| *Mimosa pithecolobioides* | 66 | 0.9436 | 0.81 | 0.89 | 0.91 | 1 |
| *Mimosa pteridifolia* | 81 | 0.9744 | 0.67 | 0.85 | 0.6 | 1 |
| *Mimosa sericantha* | 49 | 0.8243 | 0.77 | 0.93 | 0.93 | 1 |
| *Mimosa verrucosa* | 137 | 0.9068 | 0.86 | 1 | 0.94 | 0.93 |
| *Neea theifera* | 142 | 0.883 | 0.72 | 0.75 | 0.58 | 1 |
| *Ouratea hexasperma* | 232 | 0.9533 | 0.82 | 1 | 0.98 | 0.99 |
| *Oxandra reticulata* | 49 | 0.8532 | 0.59 | 1 | 0.99 | 0.9 |
| *Palicourea rigida* | 357 | 0.9556 | 0.53 | 0.77 | 1 | 0.94 |
| *Paralychnophora bicolor* | 73 | 0.9368 | 0.95 | 0.87 | 0.84 | 1 |
| *Parinari obtusifolia* | 140 | 0.9146 | 0.79 | 0.89 | 0.63 | 1 |
| *Parkia platycephala* | 145 | 0.899 | 0.51 | 0.63 | 1 | 0.63 |
| *Peltogyne confertiflora* | 131 | 0.9511 | 0.64 | 0.91 | 1 | 0.9 |
| *Persea venosa* | 84 | 0.8734 | 0.57 | 0.4 | 1 | 0.32 |
| *Piptocarpha rotundifolia* | 279 | 0.8366 | 0.96 | 1 | 0.69 | 0.97 |
| *Platonia insignis* | 92 | 0.9072 | 0.56 | 0.79 | 1 | 0.75 |
| *Plenckia populnea* | 275 | 0.882 | 0.82 | 0.93 | 0.96 | 1 |
| *Pouteria ramiflora* | 365 | 0.8327 | 0.6 | 0.79 | 1 | 0.8 |
| *Protium ovatum* | 206 | 0.9052 | 0.77 | 0.95 | 0.98 | 1 |
| *Pseudobombax longiflorum* | 138 | 0.8339 | 0.75 | 0.96 | 0.91 | 1 |
| *Qualea multiflora* | 325 | 0.8535 | 0.81 | 0.84 | 0.87 | 1 |
| *Qualea parviflora* | 361 | 0.8402 | 0.28 | 1 | 0.14 | 0.74 |
| *Rauvolfia weddeliana* | 50 | 0.8365 | 0.88 | 0.95 | 0.94 | 1 |
| *Roupala montana* | 369 | 0.8303 | 0.94 | 0.98 | 1 | 0.99 |
| *Rourea induta* | 365 | 0.864 | 0.87 | 0.89 | 1 | 0.9 |
| *Sabicea brasiliensis* | 270 | 0.8922 | 0.84 | 0.95 | 1 | 0.64 |
| *Salacia crassifolia* | 130 | 0.93 | 0.42 | 0.87 | 1 | 0.69 |
| *Salacia grandifolia* | 36 | 0.9144 | 0.95 | 0.9 | 0.91 | 1 |
| *Salvertia convallariodora* | 58 | 0.8732 | 0.74 | 0.89 | 0.88 | 1 |
| *Schefflera macrocarpa* | 293 | 0.879 | 0.58 | 0.81 | 0.9 | 1 |
| *Schefflera vinosa* | 293 | 0.8603 | 0.78 | 0.87 | 0.85 | 1 |
| *Simarouba versicolor* | 241 | 0.8676 | 0.72 | 0.93 | 0.85 | 1 |
| *Strychnos pseudoquina* | 180 | 0.883 | 0.68 | 0.86 | 1 | 0.83 |
| *Stryphnodendron adstringens* | 284 | 0.8775 | 0.67 | 0.83 | 1 | 0.94 |
| *Stryphnodendron rotundifolium* | 142 | 0.9004 | 0.85 | 0.77 | 1 | 0.85 |
| *Syagrus comosa* | 119 | 0.9113 | 0.83 | 0.9 | 0.87 | 1 |
| *Syagrus flexuosa* | 157 | 0.9052 | 0.61 | 0.86 | 0.93 | 1 |
| *Syagrus oleracea* | 48 | 0.9283 | 0.49 | 1 | 0.62 | 0.69 |
| *Tachigali aurea* | 254 | 0.8487 | 1 | 0.9 | 0.9 | 0.86 |
| *Terminalia argentea* | 338 | 0.8381 | 0.81 | 0.91 | 0.83 | 1 |
| *Terminalia fagifolia* | 290 | 0.8666 | 0.74 | 0.75 | 0.74 | 1 |
| *Tocoyena formosa* | 387 | 0.8265 | 0.62 | 0.82 | 0.73 | 1 |
| *Unonopsis guatterioides* | 99 | 0.921 | 1 | 0.74 | 0.71 | 0.52 |
| *Vatairea macrocarpa* | 160 | 0.8727 | 0.74 | 0.83 | 0.81 | 1 |
| *Vellozia squamata* | 109 | 0.9496 | 0.76 | 0.34 | 1 | 0.61 |
| *Vochysia gardneri* | 148 | 0.9234 | 1 | 0.64 | 0.92 | 0.68 |
| *Vochysia rufa* | 283 | 0.8684 | 0.73 | 0.85 | 0.84 | 1 |
| *Wunderlichia crulsiana* | 25 | 0.9885 | 0.17 | 0.26 | 0.98 | 1 |
| *Wunderlichia mirabilis* | 95 | 0.9586 | 0.51 | 0.56 | 1 | 0.29 |
| *Zeyheria montana* | 289 | 0.864 | 0.67 | 0.86 | 0.87 | 1 |

**Appendix S2**: Sample point coordinates and GBIF identification used for the endemism analysis and distribution models.
